# Artificial light at night and concealment level affect detection probability of stranded seabirds: a study with 3D-printed puffin decoys

**DOI:** 10.64898/2026.10.01.756039

**Authors:** Taylor M. Brown, Lucy van Haaften, Sabina I. Wilhelm, Suzanne Dooley, Gary Burness

## Abstract

Fledglings of burrow-nesting seabirds are vulnerable to disorientation by artificial light at night (ALAN) and grounding in coastal human settlements, where most individuals presumably perish if not rescued. Accurate estimates of the effects of stranding on seabird populations remain elusive due to several levels of uncertainty, such as the number of stranded seabirds estimated to remain undiscovered by rescuers. One species subjected to stranding is *Fratercula arctica* (the Atlantic puffin). We hypothesized that the detection probability of *F. arctica* fledglings by the Puffin Patrol rescue program, operating in Newfoundland, Canada, is affected by light level, concealment level, and search effort. We deployed realistic *F. arctica* decoys in locations of varying light and concealment levels throughout 2 sites that are searched nightly by rescuers and measured the amount of time and search effort required until each decoy was found.

Approximately 78% more search effort was required to find a decoy in darkness (32.0 min) compared with illuminated conditions (18.0 min), and up to 100% more search effort was required to find a decoy placed in concealed conditions (35.5 min) than one in the open (17.5 min). After 2 hours of searching, on average over 10% of decoys were still not found, although each additional minute of search effort increased detection rate of an individual decoy by 0.6%. In combination with recent advances in our knowledge of seabird post-stranding behaviour, our study offers a first step in the development of correction factors which can be adapted to site- and species-specific requirements of other rescue programs around the world that rely on volunteers to search for stranded birds. Application of such correction factors will hopefully result in better estimation of mortality rates and their resultant effects on populations.

**LAY SUMMARY:**

- Each year when they leave their nests, young *Fratercula arctica* and certain other seabirds become stranded in coastal towns, cannot re-orient to the sea, and are rescued by volunteers
- It is unknown how many birds remain unfound each year, but this information is critical to estimate effects of stranding on populations
- We deployed 3D-printed *F. arctica* decoys at real stranding locations in Newfoundland to determine if light levels, concealment levels, and/or search effort affect decoy detection by volunteers
- In general, volunteers found decoys more easily in locations of brighter light and lower concealment, but still missed on average ≥10% of decoys after 2 hours regardless of search effort
- Our results offer a first glimpse into the proportions of stranded seabirds potentially never found by rescue programs and thereby provide a starting point for estimating the true magnitude of stranding and its effects on populations

## INTRODUCTION

The presence of artificial light at night (ALAN) has been linked to the dysregulation of biological systems in myriad species (Rich and Longcore 2006; Gaston et al. 2021; Jägerbrand and Spoelstra 2023), and between 1992 and 2017 (the most recent timespan over which it was estimated) its emittance increased by at least 49% globally (Sánchez de Miguel et al. 2021). One way in which ALAN affects animal behaviour specifically is by causing disorientation and disruption of natural movement patterns, leading to exhaustion, collision with anthropogenic structures, stranding, and/or predation (Rich and Longcore 2006; Marangoni et al. 2022).

More than 70 seabird species worldwide exhibit nocturnal stranding (or “grounding”), with individuals usually found in artificially-illuminated spaces (Gilmour et al. 2023). Most are burrow-nesting seabirds of order Procellariiformes, but *Fratercula arctica* (Atlantic Puffin) of order Charadriiformes are also impacted (Wilhelm et al. 2021; Brown et al. 2024). *F. arctica* strand in human settlements near their colonies in Europe and North America, with ALAN identified as the likely causal factor (Rodríguez et al. 2017; Brown et al. 2024; S. Hammer, pers. comm., Faroese Environment Agency). Across species, fledglings on maiden voyages from their nests appear to be most susceptible to becoming stranded (Rodríguez et al. 2017). Grounded fledglings are rarely able to regain flight in anthropogenic landscapes, and are therefore highly susceptible to mortality by predation, vehicle collisions, and dehydration despite generally high initial survival rates (e.g., Telfer et al. 1987; Ainley et al. 2001; Miles et al. 2010; Rodríguez et al. 2012). Many affected seabird species are globally imperiled and already face compounding threats (Croxall et al. 2012; IUCN 2024), but additional mortality due to ALAN-induced grounding has been identified as having further negative effects on population growth in at least 3 species (Ainley et al. 2001; Fontaine et al. 2011; Gineste et al. 2016).

In most locales affected by seabird strandings, there are no standardized survey programs in place to estimate the number of birds grounded by ALAN. Instead, non-standardized nocturnal rescue programs that rely on public participation allow for at least a crude approximation of the magnitude of stranding-related mortality, but accurate estimation is hindered by various levels of uncertainty, including the total number of fledglings produced by a population, the number of fledglings that become grounded by lights, and the number of stranded fledglings that go undiscovered or unreported (Rodríguez et al. 2017). Detection probability estimates can vary widely on a landscape scale, from as high as 100% to less than 50%, but remain an understudied factor in calculating the magnitude of seabird fallout (Vorsino et al. 2023). Additionally, deceased stranded fledglings are commonly underreported to rescue programs, leading to both detection rates and estimated mortality rates that are artificially lowered (Ainley et al. 2001). Rescue programs universally find more birds in illuminated compared to dark areas (Rodríguez et al. 2017), but it is unclear to what extent this pattern is driven by seabird versus rescuer behaviour. If rescuers are biased toward searching or more efficient at finding birds in illuminated areas, or both (Brown et al. 2023), then a considerable proportion of stranded individuals may remain undiscovered. Thus, an estimation of seabird detection probability across light as well as concealment levels would be useful in the global effort to accurately quantify ALAN-related seabird stranding and mortality rates.

In the province of Newfoundland and Labrador, Canada, the Witless Bay Ecological Reserve protects a group of islands hosting close to 400,000 breeding pairs of *F. arctica*, the largest breeding concentration of the species in North America (Lowther et al. 2020; Iles et al. 2025). In August of each year, fledgling *F. arctica* (hereafter “pufflings”) leave their nests for the first time and several hundred (at minimum 0.2% of the fledgling cohort) are found stranded along an approximate 15-km section of coastline near the reserve annually (Wilhelm et al. 2021). Since 2004, the volunteer-based “Puffin Patrol” rescue program has conducted nightly searches for stranded pufflings along the affected section of coastline and has identified several distinct “hotspots” where stranded pufflings are reliably found. These hotspots are generally well-illuminated areas of human infrastructure along the coastline, within ∼4 km of the main *F. arctica* colonies in the reserve (Wilhelm et al. 2021; Brown et al. 2024). Volunteers (especially those with children) eager to find a puffling are often informed by organizers of the reliability of these hotspots; also, volunteers who have found pufflings at these hotspots previously are presumably more likely to search there. Both of these circumstances may drive a bias toward increased search effort and increased numbers of pufflings found in those areas (Brown et al. 2023).

We hypothesized that 3 extrinsic factors affect detection probability of stranded pufflings by the Puffin Patrol: illumination level, concealment level, and search effort. To eliminate potential biases in detection probability caused by intrinsic factors such as mobility, behaviour, and variation in body size of live birds, and to control for the possible effects of predation and number of pufflings available to be found, we used realistically-sized and -coloured 3D-printed puffling decoys for this experiment. We placed decoys in dark, dimly illuminated, and brightly illuminated locations which also varied in level of concealment, throughout the 2 hotspots visited regularly by rescue volunteers. We chose to estimate detection probability at these 2 heavily searched sites rather than across the entire fallout zone, where it would be nearly impossible to estimate search effort or detection times. By monitoring the sites in real time, we could measure search effort and the time to detection of each decoy with relatively high precision. We predicted that detection would increase with illumination level, decrease with concealment level, and increase with search effort; and that more search effort would be required to find decoys in low illumination or high concealment. To visualize the point of diminishing returns on search effort, we also modeled the relationship between search effort expended and proportion of decoys found.

## METHODS

### Materials

We printed 60 life-size, three-dimensional (3D) puffling decoys following an initial 3D file design (by SaviMade; savimade.ca) created using photographs and measurements of living *F. arctica* fledglings (see Supplemental Material). Each decoy was 18 cm tall × 10.5 cm wide × 23 cm from beak tip to tail tip and was painted to mimic real pufflings (Figure 1). Although we only needed 20 decoys for our experiment, we created 60 to account for potential loss, damage, and theft.

**Figure 1.**
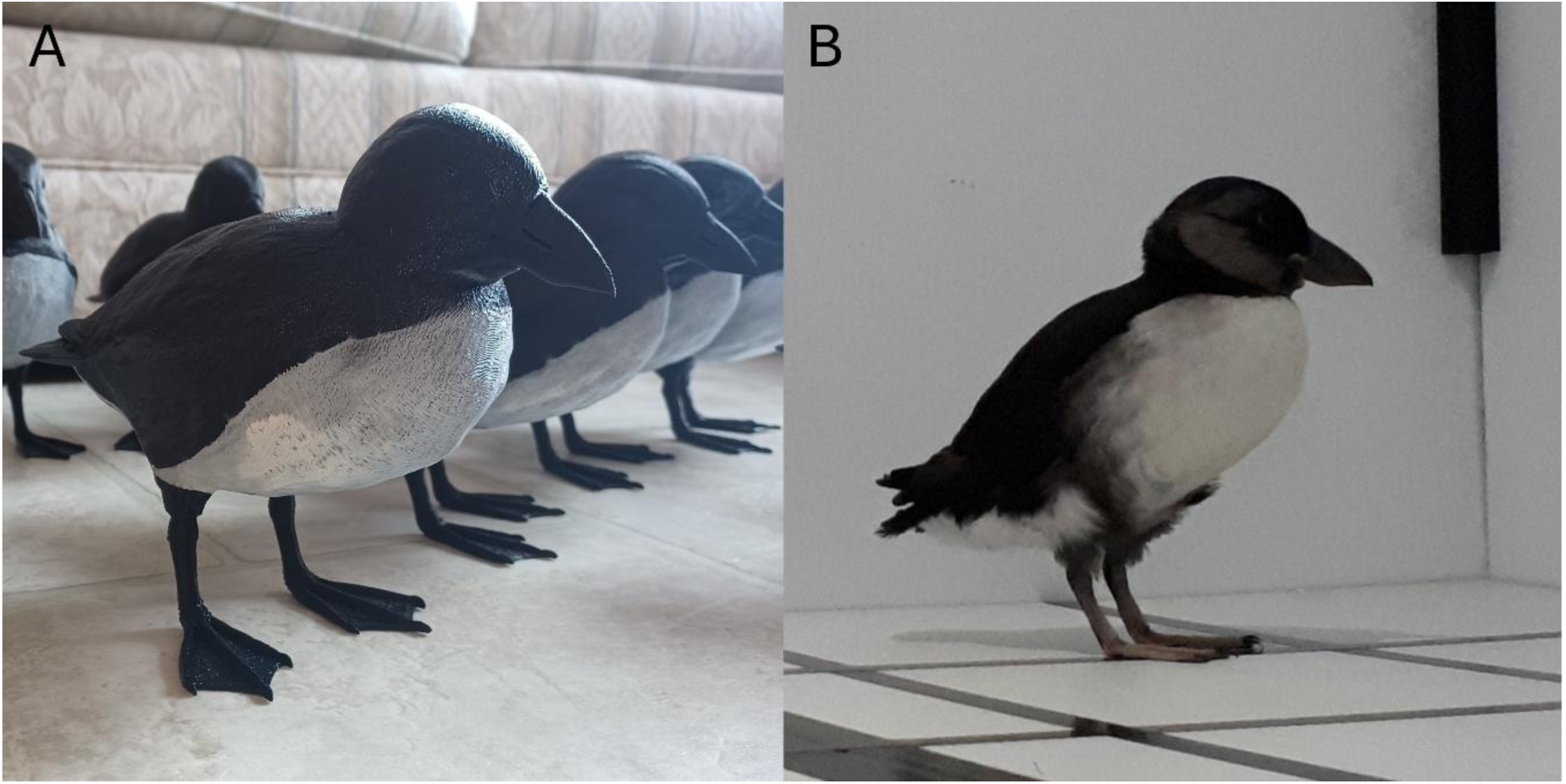
A) Several puffling decoys all assembled, painted, and ready to be deployed. B) A living puffling, used as a visual reference for the creation of puffling decoys.

### Study Sites

We conducted our experiment on 12 nights between 6-27 August 2023, at 2 sites in Witless Bay, Newfoundland and Labrador, Canada: a small commercial building that included a convenience store (hereafter “convenience store”; 47.279272°N, 52.831533°W) and the BGI crab plant (hereafter “crab plant”; 47.279404°N, 52.823854°W; Figure 2; Table 1). We chose these sites as they are stranding “hotspots” where rescue volunteers can reliably find stranded pufflings, and as such are 2 of the most heavily searched areas. The likeliest source of most, if not all, pufflings stranded in Witless Bay annually is the colony on Gull Island, located ∼3.8 km from the crab plant and ∼4.3 km from the convenience store at its closest point. We conducted the experiment on the properties surrounding each of the 2 buildings and delineated the boundaries of the total area to ensure that the area searched for the experiment at each site was approximately equal (average of 5331 m^2^; Figure 2).

**Figure 2.**
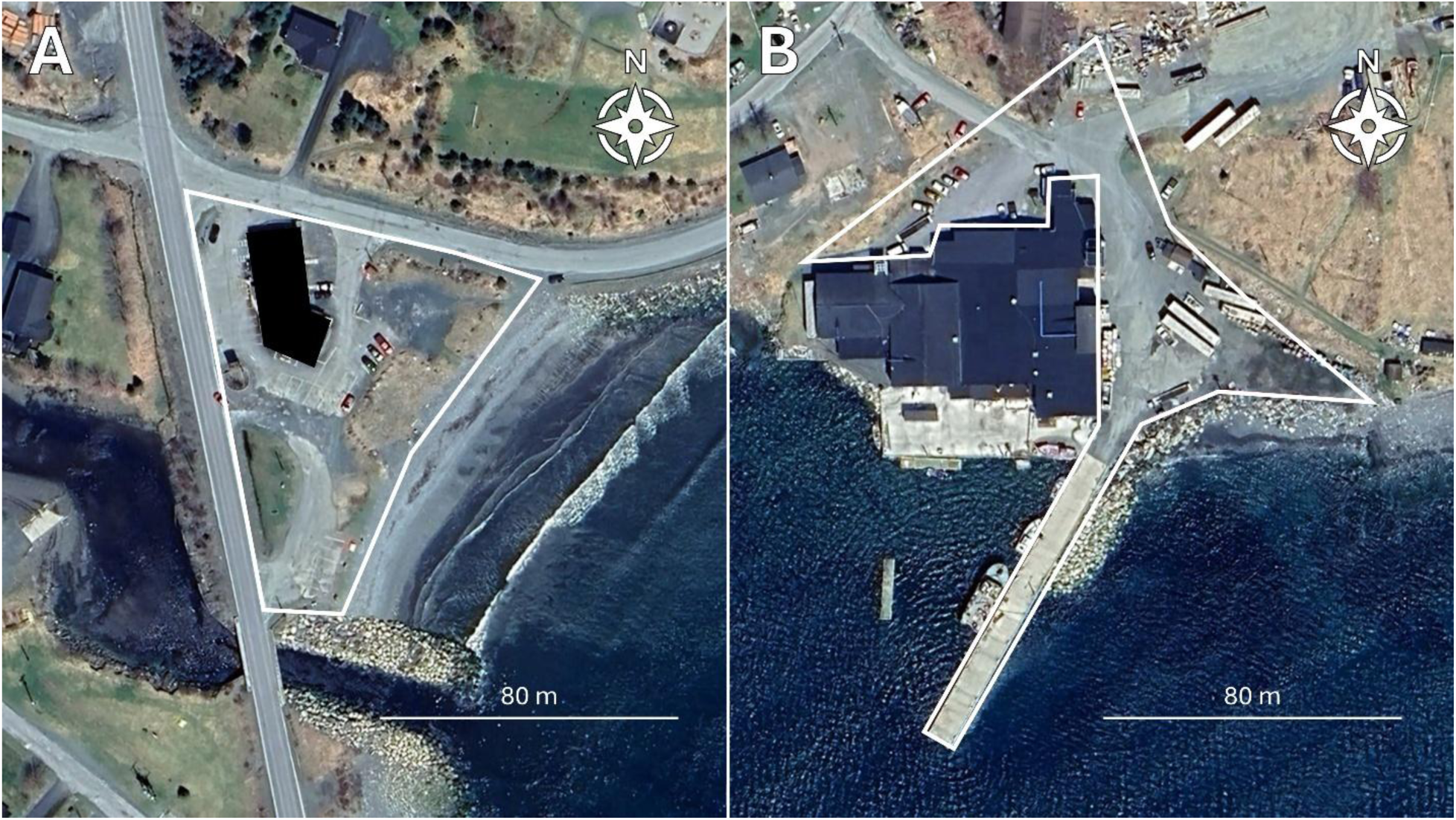
Satellite imagery (courtesy of Google Earth) of the 2 sites used in this experiment, with associated white perimeters demarcating the boundaries used for decoy deployment. A) Convenience store (note the black box, denoting the area of the building which was excluded from area calculations). B) Crab plant. The search area of the convenience store site was 5265 m^2^ (total area = 5709 m^2^ minus the area occupied by the building on site); the crab plant was only accessible on 2 sides of the building, and the search area was estimated at 5396 m^2^.

**Table 1.**
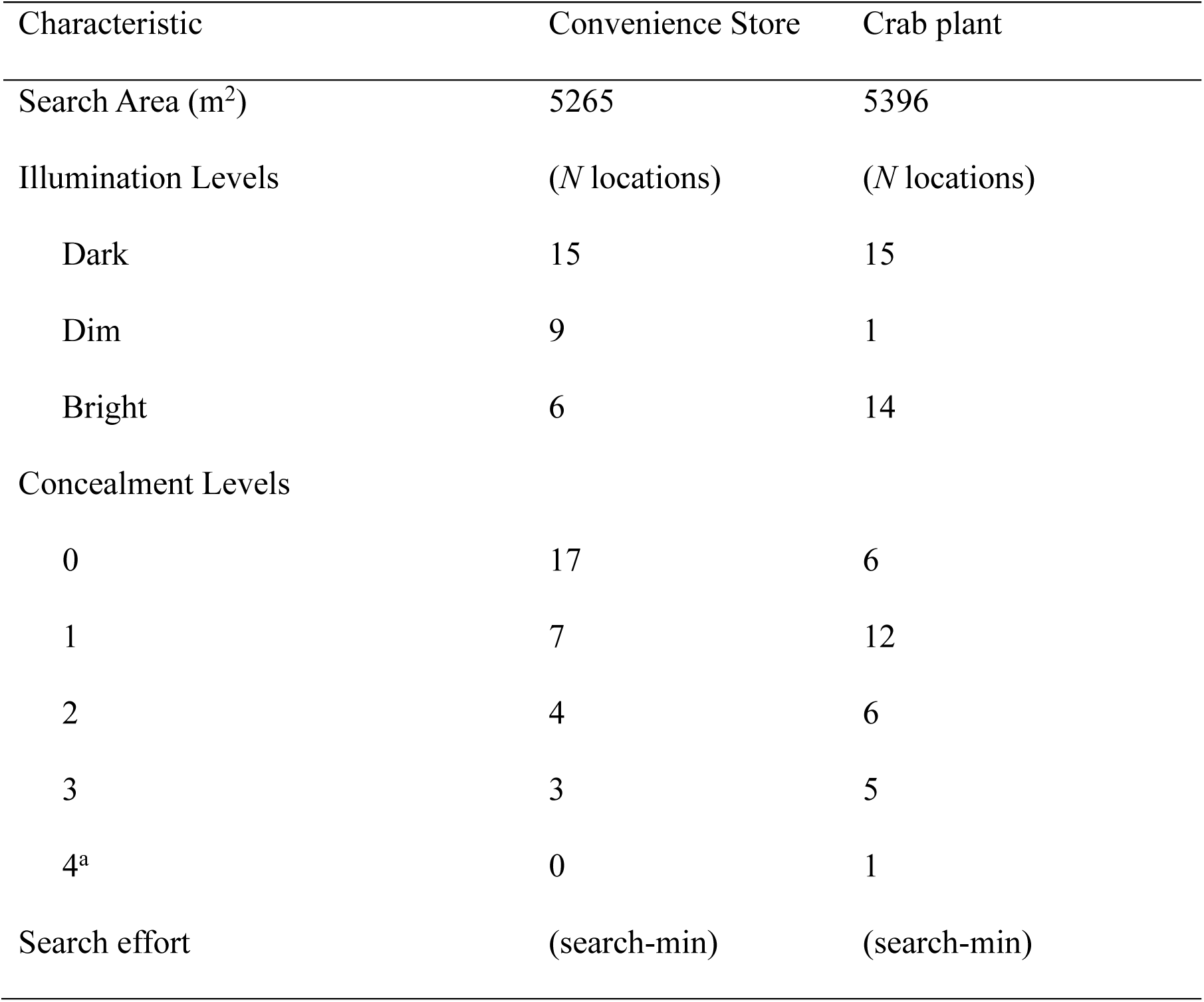
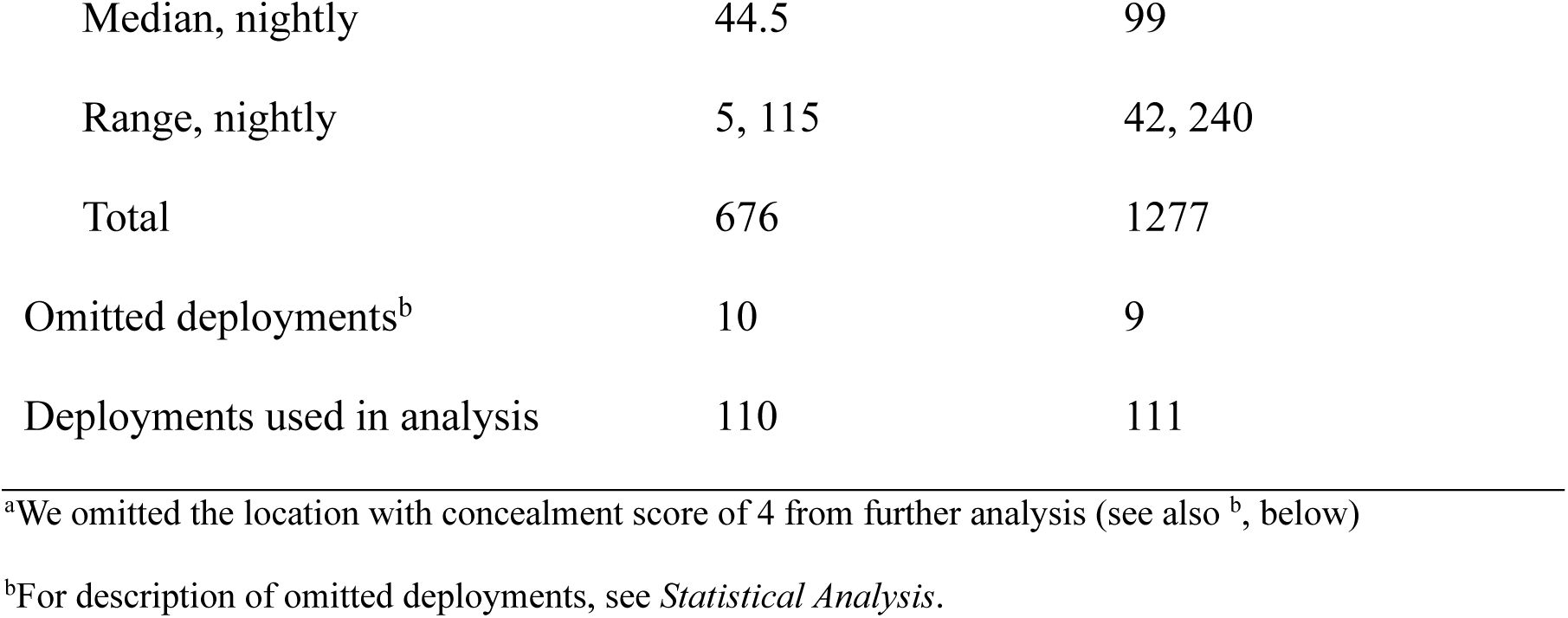
Characteristics of each of the 2 sites where puffling decoys were deployed.

### Location Selection

We selected 15 illuminated locations and 15 dark locations at each of the 2 sites using the unaided eye (i.e., without instruments) for the placement of puffling decoys (60 locations total) and marked their coordinates with a GPS unit (Garmin GPSMAP 64s or 64sx; Olathe, KS, USA; Table S1). After the experiment was complete (on 28 August 2023; a relatively clear night with the moon 91% illuminated; moongiant.com) we measured light levels in units of lux at all 60 locations with a spectrometer (SRI-2000 Spectral Light Meter; Allied Scientific Pro; Gatineau, Quebec, Canada) held ∼20 cm above the ground (the approximate height of a decoy). The spectrometer could not detect illuminance readings below 10 lux (approximately the illuminance perceived by the human eye during civil twilight; AMS 2025) and returned these readings as 0 lux. The spectrometer was therefore not accurate enough to quantify real differences between dark and dimly illuminated locations. As a result, we subclassified illuminated locations into “dim” (<10 lux; reads as “0” on the spectrometer; see above) and “bright” (≥10 lux; Table 1). “Dark” locations were defined as both subjectively dark in appearance during the initial location selection process, and those which were read by the spectrometer as 0 lux at 20 cm above ground level. These designations made sense in a real-world context because, for example, directly beneath a typical (high pressure sodium type) streetlight within the convenience store study site we measured a maximum irradiance of 14 lux at 20 cm above the ground (location B05; Table S1). We used these 3 light level categories (dark, dim, and bright) in statistical analysis and include illuminance readings in Table S1 for information purposes only. It was not possible to equalize the number of “dim” and “bright” locations between the 2 sites due to differences in site lighting and layouts, but we accounted for this by stratifying by site in our analysis (see *Statistical Analysis*, below). Although we acknowledge that the use of spectrometer-quantified illuminance measures in our analysis would be ideal, the inclusion of such measurements would not change the results, as our 3 categories were based on subjective human perception of light level (as are units of lux).

To mimic true stranding behaviour of live pufflings, at least in terms of where they are found by rescuers, we only selected locations on the ground (i.e., never on top of objects) and selected locations that varied in levels of illumination and concealment (Table S1). Across locations we varied the level of concealment on a scale based on an imaginary cube of 1 m^3^ surrounding the decoy’s precise location, not including the ground: e.g., if the decoy was concealed by a wall or other object to within 1 m on only one side, the decoy received a concealment score of 1 (Figure S1). Of the 60 locations, most had concealment scores of 0-3 (*n* = 59). We had a single location with a concealment score 4 (with 4 deployments thereof), but we omitted this location from statistical analyses as there were no replicates (Table 1). Due to the heterogeneity of the 2 sites in terms of topography, obstacles, and business activities it was not possible to perfectly equalize the number of locations with each level of concealment between the 2 sites, although the best effort was made to do so.

### Participants

When volunteers arrived at the Puffin Patrol headquarters in Bay Bulls (47.315178°N, 52.813887°W) at 2200 hours on the nights of this experiment, they were provided information about the experiment and asked if they would like to participate (for details, see *Ethical Note* and Supplemental Material). Fifty-five volunteers participated in the experiment by signing consent agreements and we allowed them to participate on as many nights as they wished, without signing additional agreement forms. They are hereafter referred to as anonymous “Participants” in those contexts pertaining to decoy detection specifically. Otherwise, we refer to volunteers more generally because in the field it was not always possible to guess, and we as the Experimenters were not permitted to ask, a volunteer’s participatory status in this experiment. All volunteers (Participants and non-Participants alike) searched for real pufflings and were counted toward metrics of search effort in this experiment, but only Participants were responsible for collecting decoys. To be clear, Participants never received any indication as to how many decoys were available to be found, nor where they may be found (see Supplemental Material).

### Experimental Procedure

Each night, we (the “Experimenters”) deployed 20 decoys, 10 at each site. Of the 10 decoys at each site, 5 were placed in dark locations and 5 in illuminated locations (dim and bright combined; separated during analysis). Because there were 30 possible locations at each site (prior to omission of one location at the analysis stage), we randomly determined which 10 locations we would use at each site on each night of the experiment, sampling without replacement over 3-day intervals. That is, all locations at each site were used over a 3-day period before randomly re-sampling. Because the experiment spanned 12 nights and we deployed 20 decoys per night, we used each location 4 times (total of 240 deployments). Despite using each individual decoy location 4 times throughout the experiment, we treated each deployment as an independent sub-sample for several reasons. First, precision of deployment location varied by up to several metres due to imperfect GPS locational accuracy and intra- and inter-Experimenter variability in decoy placement (when this variation resulted in a decoy receiving a different category of light or concealment level than intended, we omitted this particular deployment from subsequent analysis; see Statistical Analysis, below, and Table 1). Second, the use of each deployment location was randomized over 3-day intervals without replacement. This was important because most Participants were tourists/visitors who would have searched for decoys on a single night only or, if on more than one night, would have searched on consecutive nights. As a result, most Participants would have encountered a decoy at a specific deployment location only once. Third, our method of randomization of deployment locations meant that even if a Participant took part in the experiment on multiple nights, they were never exposed to decoys placed in the exact same complement of locations more than once. Furthermore, if decoys had already been detected by other Participants and removed from the experiment for the remainder of a given evening, the locations where remaining decoys could be encountered (and number of decoys that could be encountered) by any subsequent Participant would differ even more.

Between 2130-2200 hours on each night of the experiment, the 2 Experimenters (one stationed at each of the 2 sites) deployed the 10 decoys allocated to that site in time for the experiment to begin at ∼2200 hours (range in start time 2143-2159 hours). The experiment began whenever the last decoy was deployed at each site; as such, the start time was allowed to vary slightly between sites (median variation in start time between the 2 sites on the same night was 6 minutes; maximum 12 minutes). The start time of 2200 hours coincided both with complete darkness and with the beginning of the second “shift” of Puffin Patrol volunteers (2200-2400 hours). To reduce any potential bias in the direction of decoy placement and resulting differences in visibility, we randomly assigned to each decoy on each night one of the 4 main cardinal directions (north, east, south, and west), toward which it would be faced. During deployment, Experimenters used GPS units both to locate the designated locations where decoys were to be placed and to ensure the decoys faced their assigned cardinal directions. Following decoy deployment, Experimenters stationed themselves at a designated location in each site, chosen to maximize the visibility of site entrances/exits.

As an index of search effort for each site on each night, Experimenters recorded when and how many volunteers arrived and departed from each site. Volunteers were only considered to be present at the site if they were seen wearing reflective vests (issued by the Puffin Patrol) and searching on foot within the site boundary. Arrival time of volunteers was recorded as the time at which they were first seen on foot within the site boundary. Because all volunteers were included in estimates of search effort, this metric is thus inflated as compared with the search effort of Participants alone, but in a way that is consistent across both sites and across all illumination and concealment levels. We assumed that search effort of Participants alone (which was unquantified in this study) and search effort by volunteers in general (which is reported in this study) were highly correlated.

We quantified search effort for each site on each experimental night in units of “search-minutes” (i.e., one search-minute is equivalent to one volunteer searching for one minute). For example, if there were no volunteers present for 80 minutes of the experiment (0 × 80 = 0), then 5 volunteers were present for 10 minutes (5 × 10 = 50), and then 2 volunteers were present for 30 minutes (2 × 30 = 60), the search effort calculated for that site on that night would be 0 + 50 + 60 = 110 search-minutes. Note that we were not concerned about equivalency in search effort among different group sizes; we assumed that, e.g., 2 volunteers searching for 10 minutes was equivalent to 10 volunteers searching for 2 minutes (i.e., both equal 20 search-minutes).

Experimenters recorded the time at which each decoy was found. Experimenters had planned to also record the time and number of real pufflings found by volunteers. However, because of mass chick mortality at the Gull Island colony in 2023 (S.I.W, pers. obs.) there were few pufflings grounded overall and zero pufflings found grounded at either of the 2 experiment sites on our experimental nights. Occasionally, Participants collected multiple decoys while out of view of the Experimenter, and it was only possible to assign a single “time found” to all these decoys, which was the time that they were given to the Experimenter. The experiment ended at a given site when all decoys at that site were found, or when all volunteers had returned to Puffin Patrol headquarters to sign out, or at 2400 hours, whichever occurred first (median experiment duration, 117.5 minutes; range 22-132 minutes). Experimenters collected any undetected decoys at the end of each night.

### Statistical Analysis

Due to external circumstances beyond our control (e.g., lighting changes at the businesses we used as our study sites, decoys stolen) and occasional miscommunication between Experimenters regarding decoy deployment, there were 19 deployments distributed across all 12 nights of experiments which were excluded from analysis (Table 1), leaving a final sample size of *n* = 221 deployments. Of these, 110 were in darkness and 111 were in light (38 in dim light; 73 in bright light). In terms of concealment level, 80 deployments were in concealment = 0, 76 deployments in concealment = 1, 36 deployments in concealment = 2, and 29 deployments in concealment = 3. All statistical analyses were conducted in program R (ver. 4.3.0; R Core Team 2023), and we report results with exact *P* values (*α* = 0.05 was used to denote statistical significance).

### Proportions of decoys found in each level of illumination and concealment

First, we calculated the percentages of decoys found in each of the 3 light levels (dark, dim, and bright) and separately in each of the 4 concealment levels (0, 1, 2, 3). We then conducted chi-square tests for homogeneity to determine if the proportions of detected versus undetected decoys differed among levels of each of these 2 variables (without controlling for or holding constant any other factors).

### Decoy detection rate relative to levels of illumination, concealment, and search effort

For the purpose of our analysis, detection rate was calculated relative to regular elapsed time, which includes time spans in which there were no volunteers present. We conducted this analysis to understand, on an average night when the Puffin Patrol is operational, how our variables of interest affect the detection rate of decoys. This allowed us to estimate how long (in elapsed time) real pufflings may await rescue (visualized as persistence probability over time; see Supplemental Material) across different levels of illumination, concealment, and search effort, with the effect of each covariate assessed while adjusting for those of the others. We first determined the “fate” of each deployed decoy as either detected (“1”) or undetected (“0”). We then calculated the “fate time” of each decoy as the number of minutes that elapsed until it was found, using the time at which all decoys had been deployed at a given site as the reference start time. If the decoy remained undetected until the end of the experiment, its “fate time” was the duration of the experiment (in minutes). Any decoys that remained undetected at the end of an experiment with a duration <120 minutes (no less than 116 minutes) were “censored”; our analysis contained 22 such censored observations. Censored individuals were dropped from the model as having persisted at the time that they became censored and thus did not affect estimates of detection rate beyond that time.

We tested for effects of light level (ordinal: dark [0], dim [1], or bright [2]; at the level of the individual decoy location), concealment level (ordinal: 0, 1, 2, or 3; at the level of the individual decoy location), and search effort (continuous, numeric: in search-minutes; at the level of each site on each night) on “detection rate” using Cox proportional hazards (CPH) models (Cox 1972) with the *survival* (3.6-4; Therneau 2024) and *survminer* (0.4.9; Kassambara et al. 2021) packages. “Detection rate” was defined as the instantaneous likelihood of a detection event occurring at any given time and was encoded by the CPH model incorporating both the “fates” and “fate times” of decoys. We calculated the magnitude of difference in detection rate between individual levels of a variable using the hazard ratio (“HR”) provided in the CPH model output.

Search effort differed between the 2 sites (*W* = 31.5, *P* = 0.02; Table 1). We accounted for these differences between the 2 sites in our CPH model both by testing for the effect of search effort directly, as a covariate, and by stratifying the data according to site (convenience store and crab plant) using the *strata()* function. Stratification allowed the baseline hazard functions to differ between sites while still assuming a common effect of the chosen covariates.

We tested the proportional hazards assumption using the *cox.zph()* function (package *survival*, 3.6-4) and found no violations (light level χ^2^ = 0.14, *P* = 0.93; concealment χ^2^ = 2.06, *P* = 0.56; search effort χ^2^ = 0.05, *P* = 0.83; global χ^2^ = 2.27, *P* = 0.89). We checked for multicollinearity by calculating variance inflation factors using the *vif()* function in package *car* (3.1-2; Fox and Weisberg 2019) and detected no collinearity among our 3 covariates (light level, concealment, and search effort), as their GVIF^(1÷(2×df))^ values were all between 1.00-1.02.

### Differences in search effort required to find decoys across illumination and concealment levels

We assessed whether the amount of search effort required to find an individual decoy (*y*) differed among our 3 light levels (*x*) or, separately, among our 4 concealment levels (*x*). Because search effort data were severely non-normal, we compared medians using non-parametric statistics. Specifically, we performed 2 separate Kruskal-Wallis H tests (package “*stats*”, 4.3.0) and subsequent post hoc Dunn tests (package “*FSA*”, 0.9.4; Ogle et al. 2023) for multiple comparisons. Note that concealment level was not controlled for or held constant in the comparisons among light levels, or vice versa.

### Relationship between search effort and proportion of decoys found each night

In our final analysis, we wanted to explore the shape of the relationship between search effort and proportion of decoys found, irrespective of illumination and concealment levels, so that we could estimate the point of diminishing returns. First, we calculated for each detected decoy (*n* = 163): (1) the search effort (in search-minutes) expended to find it; and (2) the cumulative proportion of decoys found that it represented on a per-site, per-night basis (e.g., the sixth decoy found out of 10 deployed decoys would be assigned a cumulative proportion value of 0.6). In the event that 2 or more decoys were found at the same time, the cumulative proportion values assigned to those decoys were tied (e.g., if the first 2 decoys out of 10 were detected simultaneously, both were assigned the value of 0.2). This cumulative proportion calculation excluded from the denominator any decoys that were omitted from analysis (described previously), with the smallest denominator being 8. We modeled the relationship between search effort and proportion of decoys found using a quasibinomial generalized additive mixed model (GAMM) with logit link function, coding “site” as a random effect, using the *gamm* function in package *mgcv* (1.9-1; Wood 2023). The model was then visualized using the functions *gratia* (0.10.0; Simpson 2024) and *ggplot2* (3.5.1; Wickham 2016).

## RESULTS

### Proportions of Decoys Found in Each Level of Illumination and Concealment

Of 221 deployed decoys, 163 (74%) were found by Participants (Table 2). Seventy-three decoys were found at the convenience store out of 110 deployed (66% detected), versus 90 found out of 111 deployed at the crab plant (81% detected). Separated according to light level, Participants found 66% of decoys deployed in “dark” locations, 76% of decoys in “dim” locations, and 84% of decoys in “bright” locations (Table 2); these differences were statistically significant (χ^2^_2, 221_ = 6.86, *P* = 0.03). When “dim” and “bright” deployments were considered together as simply “illuminated” locations, 81% of decoys were found (*n* = 90 of 111; Table 2), suggesting that even when there is some light, close to 20% of decoys still remain undetected after 2 hours.

**Table 2.** Number of decoys deployed and found in each light level, along with calculated percentage of decoys found.

|  | **Dark** | **Dim** | **Bright** | **Total** |
|---|---|---|---|---|
| Deployed | 110 | 38 | 73 | **221** |
| Found | 73 (66%) | 29 (76%) | 61 (84%) | **163 (74%)** |

Separated according to concealment level, Participants found 80% (*n* = 64 of 80) of decoys placed in the open (concealment score = 0), and this did not differ from when decoys were concealed on 1, 2, or 3 sides (χ^2^ = 5.33, df = 3 and 221, *P* = 0.15; Table 3). When deployments with concealment scores of 1-3 are considered together as simply “concealed” locations, 70% of decoys were found (*n* = 99 of 141; Table 3).

**Table 3.** Number of decoys deployed and found in each concealment level, along with calculated percentage of decoys found.

|  | **Concealment Level** |  |  |  |  |
|---|---|---|---|---|---|
|  | **0** | **1** | **2** | **3** | **Total** |
| Deployed | 80 | 76 | 36 | 29 | **221** |
| Found | 64 (80%) | 54 (71%) | 22 (61%) | 23 (79%) | **163 (74%)** |

### Decoy Detection Rate Relative to Levels of Illumination, Concealment, and Search Effort

According to our Cox proportional hazards model, search effort and concealment level each had significant effects on detection rate (the instantaneous likelihood of a detection event occurring at any given time) while adjusting for the effects of the other covariates. For each additional minute spent searching by one person (i.e., one “search-minute”) at a given site on a given night, detection rate increased by 0.6% (coefficient = 0.006; *P* < 0.001; Table 4). Note that for the concealment and illumination level covariates, the CPH model only provides comparisons for each level compared to the reference level (i.e., reference levels are concealment = 0 and light level = dark, respectively) and therefore does not compute pairwise comparisons among all levels (Table 4). Concealment played a significant role in predicting detection rate, but only up to a concealment score of 2. Decoys concealed on one side (concealment score = 1) had a detection rate 40% lower than those placed in the open (concealment score = 0) (coefficient = -0.506; *P* = 0.012; Table 4) and the detection rate of decoys concealed on 2 sides (concealment score = 2) was 57% lower than those placed in the open (coefficient = -0.852; *P* = 0.001; Table 4). Interestingly, concealment on 3 sides (concealment score = 3) did not result in any significant difference in detection rate compared to decoys placed in the open (*P* = 0.134; Table 4). Detection rate was higher, but not statistically significant, in brightly illuminated conditions compared to darkness (coefficient = 0.321; *P* = 0.087; Table 4) and detection rate did not differ between dark and dimly illuminated locations (*P* = 0.167; Table 4). For modeled persistence probabilities of decoys at the various levels of each covariate, see Supplemental Material.

**Table 4.** Results of a Cox proportional hazards model of decoy persistence based on light level, concealment level, and search effort as fixed effects, with data stratified by site. The hazard ratio (“HR”) refers to the magnitude of difference in detection rate from the reference level of each ordinal parameter (light level = dark; concealment = 0) to the parameter level indicated in column 1 or, in the case of search effort (a continuous parameter), the magnitude of difference in detection rate with each increase in search effort of 1 search-minute. A hazard ratio >1 indicates an increase in detection rate from the reference value by a factor of HR-1 and a hazard ratio <1 indicates a decrease in detection rate from the reference value by a factor of 1-HR. HR = 1.000 or 95% confidence intervals overlapping with 1.000 would indicate no difference in detection rate.

| **Parameter (level)** | **Coefficient** | **SE of Coefficient** | **Hazard Ratio (Exponentiated Coefficient)** | **95% CI of Hazard Ratio** | ***P* value** |
|---|---|---|---|---|---|
| Light Level (Dim) | 0.326 | 0.236 | 1.386 | 0.873, 2.201 | 0.167 |
| Light Level (Bright) | 0.321 | 0.188 | 1.378 | 0.954, 1.991 | 0.087 |
| Concealment (1) | -0.506 | 0.201 | 0.603 | 0.406, 0.894 | **0.012** |
| Concealment (2) | -0.852 | 0.268 | 0.427 | 0.253, 0.721 | **0.001** |
| Concealment (3) | -0.376 | 0.251 | 0.686 | 0.420, 1.122 | 0.134 |
| Search Effort | 0.006 | 0.002 | 1.006 | 1.003, 1.009 | **<0.001** |

### Differences in Search Effort Required to Find Decoys across Illumination and Concealment Levels

There was a significant effect of light level on the amount of search effort required to find each individual decoy (Kruskal-Wallis test: *H*_2_ = 12.621, *P* = 0.002, Figure 3). Seventy-eight percent more search effort was required to find a decoy placed in darkness (median search effort = 32 search-minutes) than in dim illumination (median 18 search-minutes) or bright illumination (median 18 search-minutes; Dunn test: *P* = 0.010 and *P* = 0.007, respectively; Figure 3). There was no difference in search effort required to find a decoy between dim and bright illumination (*P* = 0.541).

**Figure 3.**
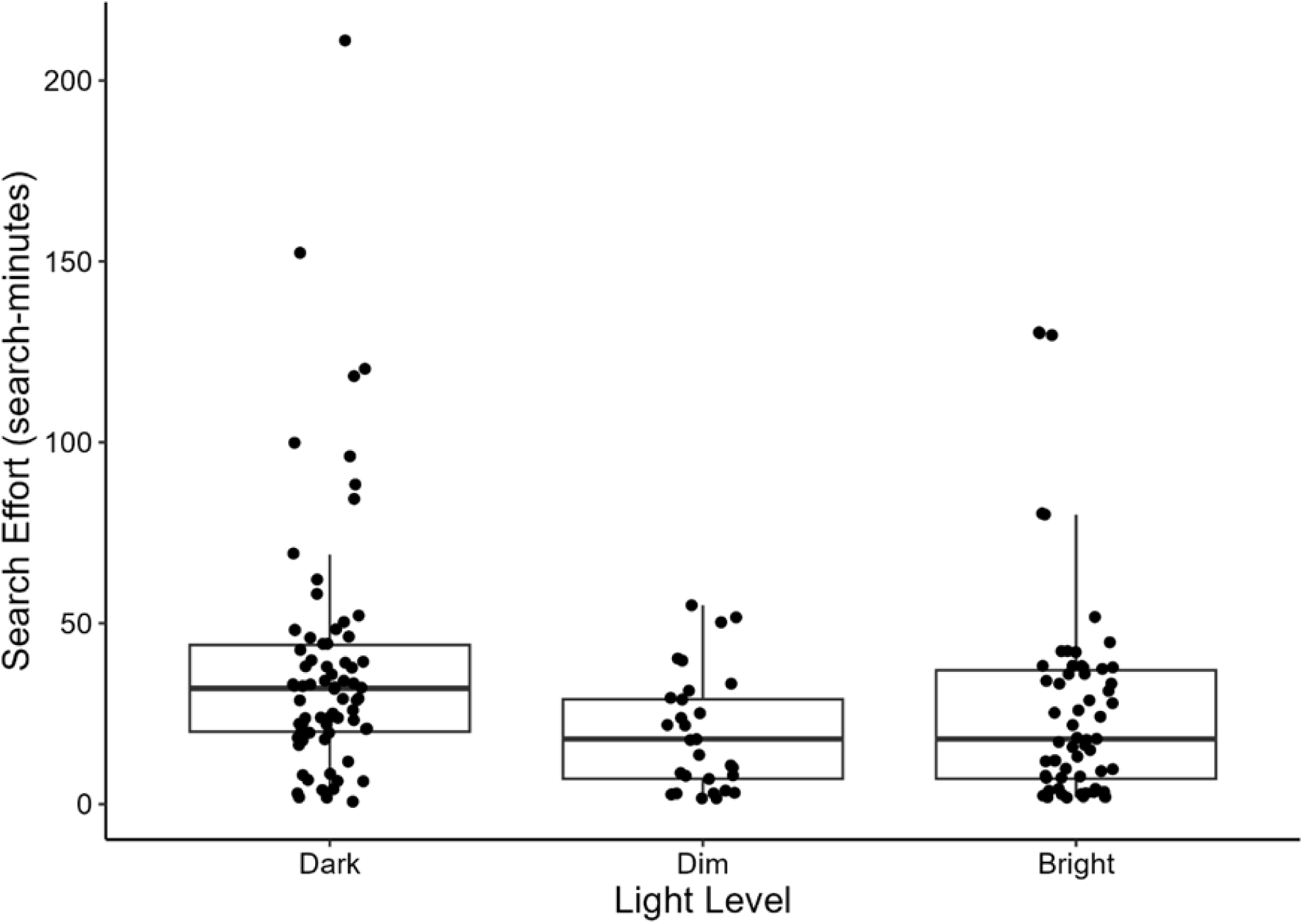
Amounts of search effort (in search-minutes) required to find each decoy (represented by individual data points) under each of 3 light levels (represented by 1 of 3 boxplots): dark (median = 32 search-minutes; *n* = 73), dim (median = 18 search-minutes; *n* = 29), and bright (median = 18 search-minutes; *n* = 61). Search effort required to find decoys in darkness was significantly higher than under dim or bright illumination; there was no difference between dim and bright.

There was also a significant effect of concealment level on the amount of search effort required to find decoys (Kruskal-Wallis test: *H*_3_ = 20.702, *P* = 0.001; Figure 4). More search effort was required when decoys were concealed on 1 side (median search effort = 32.5 search-minutes), 2 sides (median 35.5 search-minutes), and 3 sides (median 26 search-minutes) than when they were placed in the open (median 17.5 search-minutes; Dunn test: *P* = 0.001, 0.003, and 0.030, respectively; Figure 4). There were no significant differences in the amount of search effort required to find decoys among concealment levels 1, 2, or 3. It should be noted that in these comparisons we did not control for or hold constant light level.

**Figure 4.**
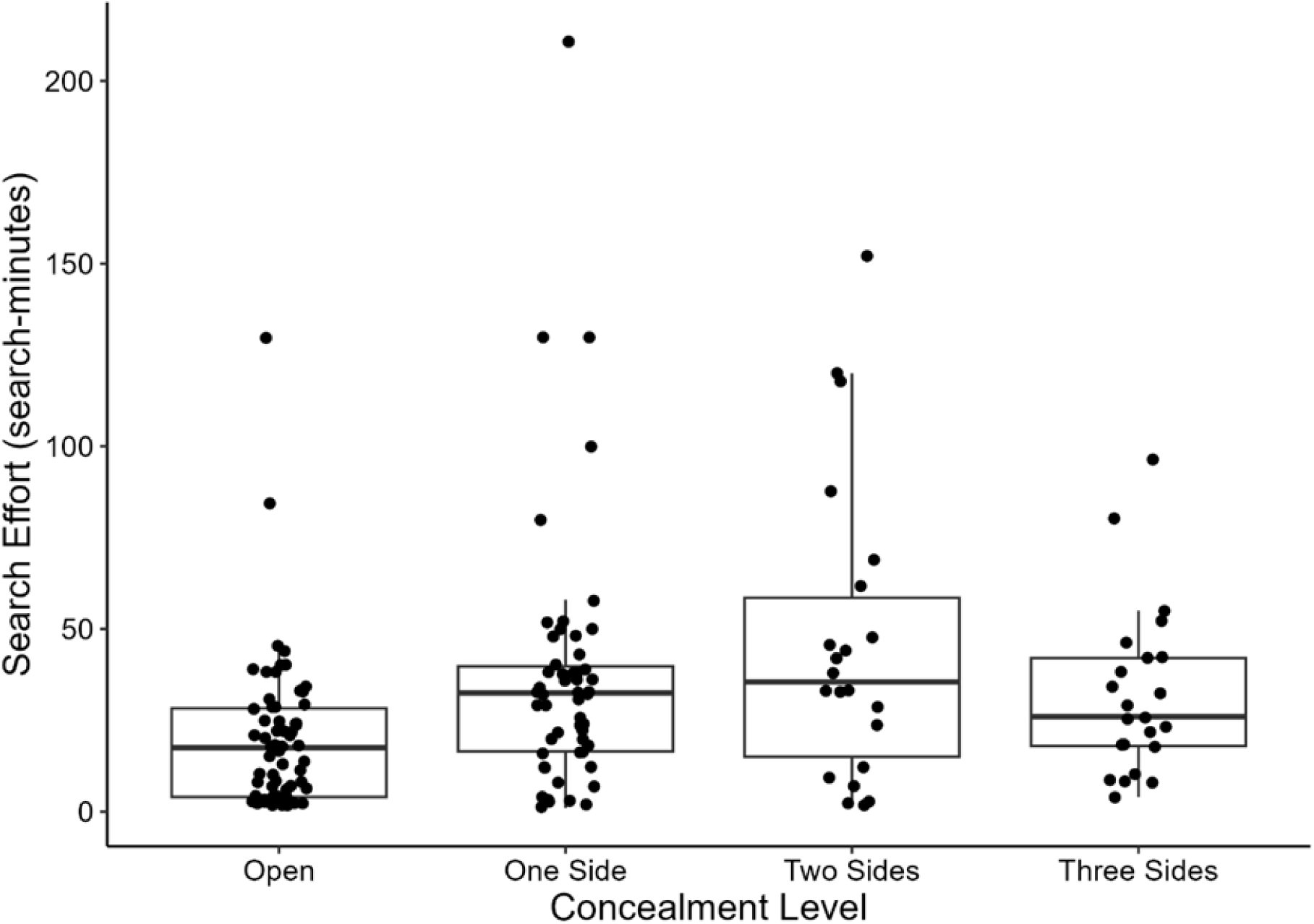
Amounts of search effort (in search-minutes) required to find each decoy (represented by individual data points) under each of 4 concealment levels (represented by 1 of 4 boxplots): those unconcealed in the open (median = 17.5 search-minutes; *n* = 64), concealed on 1 side (median = 32.5 search-minutes; *n* = 54), concealed on 2 sides (median = 35.5 search-minutes; *n* = 22), and concealed on 3 sides (median = 26 search-minutes; *n* = 23). Search effort required to find decoys was significantly less when decoys were placed in the open than when they were concealed on 1, 2, and 3 sides.

### Relationship Between Search Effort and Proportion of Decoys Found each Night

Our quasibinomial GAMM was significant in predicting the non-linear positive relationship between search effort and proportion of decoys found (effective df = 3.431, *F* = 56.350, *P* < 0.001, adjusted R^2^ = 0.601; Figure 5). Proportions of decoys found increased steeply until approximately *x* = 75 search-minutes, at which point the slope levels off just above *y* = 0.9 proportion found (Figure 5). That is, after an expenditure of ∼75 search-minutes of search effort, it is estimated that Participants would have found ∼90% of the decoys (assuming a denominator of ∼10 decoys available to be found at each site on each night, as in our experiment). The intercept of the relationship did not significantly differ from zero (estimate = 0.245, SE = 0.165, *T* = 1.484, *P* = 0.140).

**Figure 5.**
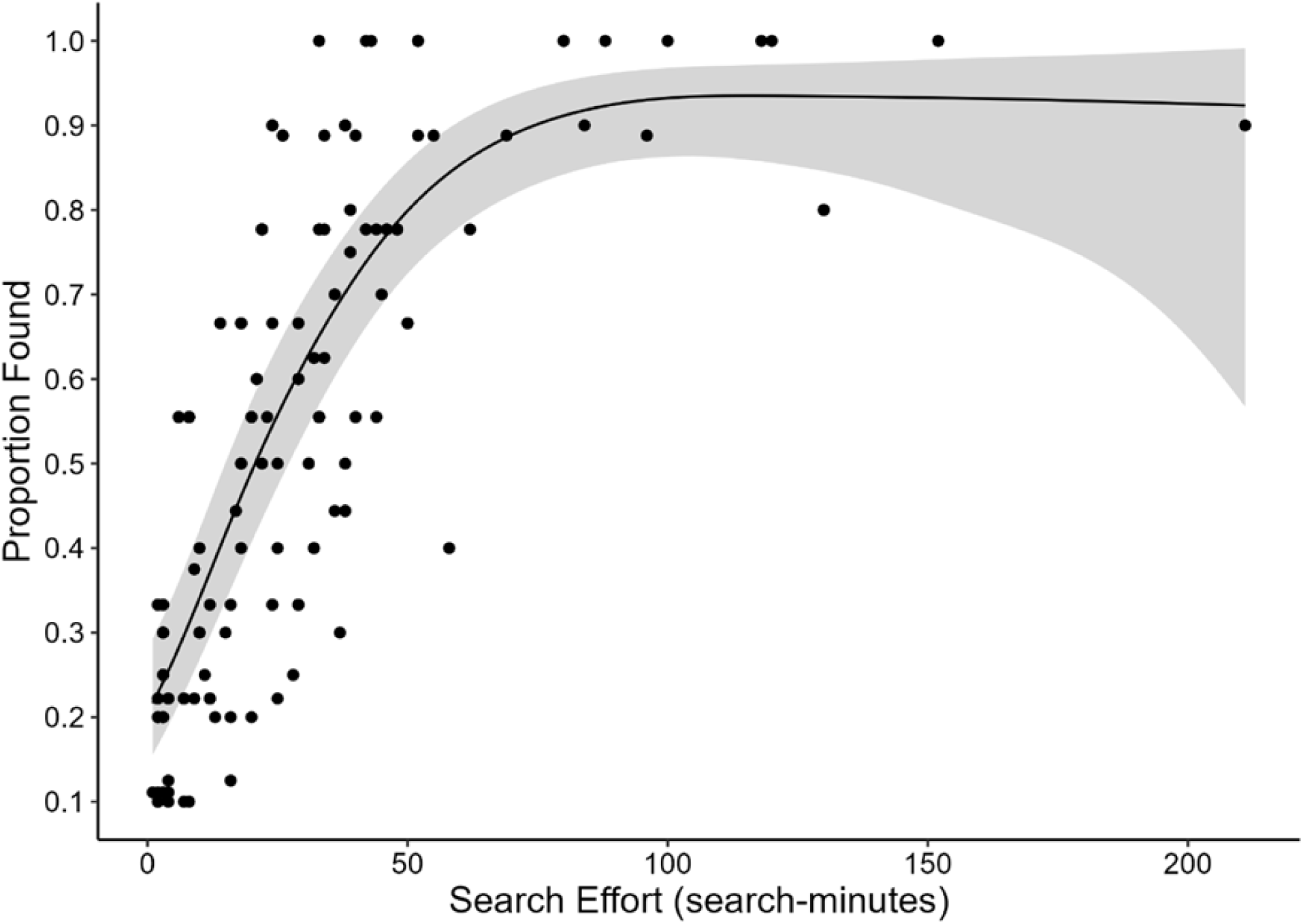
Increased search effort resulted in higher proportions of decoys found until ∼75 minutes of search effort had been expended, at which point the slope levels off just above *y* = 0.9. Individual data points represent the cumulative proportion of decoys found (*y*) at a given site on a given night, with the expenditure of *x* search-minutes of search effort. Trendline depicts generalized additive mixed model of the relationship, with associated 95% confidence intervals (shaded areas).

## DISCUSSION

Our study, as far as we know, is the first to estimate variation in detection probability of stranded seabirds by rescue program volunteers. We placed 3D-printed decoys throughout 2 puffling stranding “hotspots” over 12 nights, and measured search effort and time required to find them. We found support for our hypothesis that detection probability of stranded pufflings by the Puffin Patrol is affected by light level, concealment level, and search effort. Most of the decoys deployed at a given site on a given night were detected with the expenditure of at least 75 search-minutes of search effort, regardless of light and concealment levels. However, to our surprise, ∼10-15% of decoys still remained undetected by Participants even when in open areas and/or under bright illumination. We recognize that search effort and detection probability at these hotspots are likely higher than much of the remaining fallout zone (as also anecdotally reported for other species and in other fallout zones), and for this reason we suggest interpreting these values as maxima, while the proportions of undetected decoys should be interpreted as minima.

### Effects of Illumination and Concealment Level on Decoy Detection Probability

Higher proportions of decoys were detected with increased light levels (Table 2). However, the *rate* at which decoys were detected (i.e., the instantaneous likelihood of a detection event occurring at any given time) did not vary significantly with illumination level (Table 4; although the difference between darkness and bright light approached significance, *P* = 0.087). This lack of difference in detection rate was surprising given that an individual searcher needed to expend almost twice as much search effort to find a decoy in darkness (median 32 search-minutes) than in either of the 2 illuminated conditions (median 18 search-minutes). This could potentially be explained if Participants preferentially search in dark locations early in the search period. Participants found proportionally more decoys in illuminated locations than in dark ones (Table 2), also leading to differences in persistence probability among decoys in the 3 different light levels (Figure S2).

The relationship between concealment level and detection probability was more complex. In general, concealment on any number of sides decreased the probability that a decoy would be detected compared to when decoys were placed in the open (Table 3). Significantly more search effort was required to find decoys concealed on at least one side compared to unconcealed ones (Figure 4). Yet, surprisingly, decoys concealed on 3 sides were detected in a similar proportion (Table 3) and rate (Table 4) to those placed in the open, and slightly (not significantly) less search effort was required to find decoys concealed on 3 sides than those concealed on 1 or 2 sides (Figure 4). We acknowledge that almost all “dim” decoy locations were at the convenience store site and that this is also where search times and decoy detection rates were lower; however, we believe both of these factors are coincidental to the site layout overall rather than directly linked to one another (see Future Directions). Understanding the actual search patterns of the Participants would have been interesting, as it would allow us to better articulate the spatiotemporal relationships of where and when Participants were searching, versus where and when decoys were found. As a result, coordinators of the Puffin Patrol could provide targeted guidance to volunteers on effective search strategies.

### Implications for Live Birds and Rescue Programs

Using actual volunteers from the Puffin Patrol rescue program, our results demonstrate that rescuers show differences in detecting stranded seabirds across illumination and concealment levels. Differential detection probability is important to consider in the effort to increase the effectiveness of rescue programs, particularly in the context of post-stranding behaviour in response to light, which can vary widely across affected taxa. For example, fledgling *F. arctica* (Charadriiformes) ground at higher rates in light areas than dark ones and preferentially move toward light even after becoming stranded (Brown et al. 2024). In contrast, procellariiforms also become stranded at higher rates when illumination is higher (Rodríguez et al. 2014; Burt et al. 2024) but, once grounded, individuals remain within or move toward dark areas and even preferentially hide in concealed locations when given the opportunity (Atchoi et al. 2024; Burt et al. 2024; Brown et al. 2025). Therefore, seabird rescue efforts should employ different strategies for maximizing detection rate between e.g. charadriiforms and the much more widely-affected procellariiforms, with greater focus in the latter case on searching dark and concealed locations.

From our results, it would appear that Puffin Patrol rescuers employ relatively effective strategies for finding decoys in illuminated and open conditions, which is where we would expect most stranded pufflings to be (Brown et al. 2024). Puffin Patrol rescuers are also relatively effective at searching highly concealed, but not intermediately concealed or dark locations. This implies that they might not be quite as effective in finding stranded procellariiforms as they were at finding pufflings. That said, volunteers may unknowingly be using a relatively effective intermediate search strategy that balances search effort across illumination and concealment levels: after all, our model demonstrated they were able to find ∼90% of decoys scattered over approximately a half hectare (1.3 acres) area within a 75 minute total search-effort interval (Figure 5). Nonetheless, many additional factors may influence searcher behaviour and decoy (or seabird) detection probability, including substrate colour, general terrain, and searcher experience, among others. We controlled for these factors as best we could, but we acknowledge that they may influence the rate at which real birds would be found.

When considering living birds, the time until detection is arguably just as important as the proportion of individuals found, considering time is likely the main difference between a seabird found alive and one found deceased (e.g., Rodríguez et al. 2014) or not found at all (e.g., Podolsky et al. 1998). Despite the fact that our decoys were detected in statistically similar proportions across concealment levels (Table 3), when decoys were placed in locations of intermediate concealment (i.e., concealment level 2), the rate at which they were detected was approximately half that of those placed in the open (Table 4; hazard ratio = 0.427). Further, the search effort required to find decoys of intermediate concealment was up to double that of those placed in the open (Figure 4). This means that stranded birds that have managed to only partially conceal themselves may face greater predation risk than those in more fully concealed locations not only because they are more exposed, but also because volunteers are less efficient at finding them (Table 4; Figure 4). Although some procellariiforms may be successfully rescued up to 5 days after grounding (Rodríguez et al. 2015), rapid response times are important in ensuring high survival rates: significantly more stranded seabirds are found during nocturnal surveys (Rodríguez et al. 2014; Burt et al. 2024), and a higher proportion of them are found alive (Rodríguez et al. 2014), compared to when surveys are conducted the next morning. Rapid predation of stranded seabirds has been observed in Newfoundland, where morning surveys for living birds, carcasses, and individual body parts resulted in estimates of at minimum 300 individuals being predated over a one-month period, not including of course those individuals removed from the site without a trace (Burt et al. 2024). In fact, Podolsky et al. (1998) calculated that rapid removal of seabirds (in this case carcasses thereof) by scavengers could cause daily surveys to underestimate counts of stranded individuals by at least 17%. Clearly, high detection rates are essential in maximizing the survival of stranded seabirds, and our study offers insights on where efficiency may be lacking.

Across species and rescue programs, the proportion of stranded seabirds found dead or dying ranges from 3-43% (Rodríguez et al. 2017; Wilhelm et al. 2021). However, these estimates do not include the individuals that remain undetected, which probably suffer mortality rates approaching 100% given their extreme difficulty in regaining flight and returning to the ocean (Rodríguez et al. 2017). Indeed, each halving of “discovery rate” (in our terms, proportion of individuals detected) may effectively double the overall mortality rate of stranded individuals (Ainley et al. 2001). Our data suggest that despite their best efforts, volunteers will overlook >30% of stranded individuals in dark and intermediately-concealed locations and >10% of stranded individuals even in bright and open locations (Tables 2 and 3). These estimates are based on relatively small sites that were searched relatively intensely every single night and using decoys representing a relatively large body size (*F. arctica* fledglings on average weigh ∼250 g; Wilhelm et al. 2021). We expect the proportion of undetected individuals to only increase in cases where the area to be searched is larger (e.g., in the remainder of the fallout zone, outside of “hotspots”), or when the target species is smaller, such as the ∼50 gram *Hydrobates leucorhous* (Leach’s Storm-petrel; Wilhelm et al. 2021). In fact, smaller body sizes are linked to lower detection probability in carcass detection studies (Barrientos et al. 2018). Potentially making detection even more difficult, procellariiform seabirds such as *H. leucorhous* tend to move relatively little in darkness after becoming stranded (Brown et al. 2025), as opposed to *F. arctica* pufflings which are highly active in the absence of light (Brown et al. 2024). Further study of detection probabilities of ideally living stranded seabirds that includes interactive effects of body size, activity level, light level, and concealment level will be helpful in the development of accurate stranding data correction factors. These correction factors can then be modified and applied on a regional basis to account for variation in search effort across entire fallout zones (e.g., as in Vorsino et al. 2023).

### Future Directions

One of the strengths of our study was that we could evaluate the discovery of decoys by the members of the public directly participating in seabird search efforts. However, we also had little experimental control over the study sites, including the number and timing of volunteers entering and exiting the sites. Further, we could not be certain exactly how many volunteers were actual Participants in the experiment on any given night (as our ethics approval did not allow us to ask volunteers). If the search effort of Participants and that of volunteers were not correlated (which we assumed they were), our conclusions regarding the effects of search effort on seabird detection probability would be weakened, although patterns with respect to light and concealment would presumably be unaffected. In future, it would be ideal to quantify the exact numbers of volunteers and Participants each night and calculate their degree of correlation. Even better, a more tightly controlled experiment would include only Participants searching the experimental sites, rather than having to assume or calculate the proportionality of search effort between experiment Participants and volunteers in general.

As part of our analysis of decoy detection rate using Cox proportional hazards regression, we tested for multicollinearity among our 3 covariates of interest and found none. This is to say, the specific decoy locations we chose at our 2 “hotspot” sites appear to have varied enough in illumination and concealment in a non-correlative way to keep their effects independent for the purposes of our analysis. However, there likely exists some natural correlation (whether positive or negative, or a combination) between illumination and concealment levels across our sites and in the fallout zone at large, which we were not able to account for in this study. For example, at our chosen sites, many light sources were mounted on the walls of the buildings that also served in many instances as sources of concealment for those decoys placed next to them (i.e., a positive correlation: high concealment, high illumination), but in other cases, streetlights illuminated open parking lots and stretches of road (negative correlation: low concealment, high illumination) or, indeed, roads in areas of lower habitation may not have streetlights at all (positive correlation: low concealment, low illumination). Layout and relative accessibility of a site may additionally influence search patterns and effort, the effects of which may be difficult to disentangle from those site-specific differences in illumination and concealment discussed above (e.g., if a site is relatively open and illuminated, searchers may deem their search effort to be sufficiently thorough with only a drive through the site or a superficial scan). A thorough assessment of the relative contributions of, and potential for correlations between, illumination, concealment, site layout, and search effort was beyond the scope of this study. Future studies of seabird detection probability would benefit from such an assessment in order to eventually develop stranding data correction factors that are also allowed to vary spatially.

Our study used stationary plastic decoys to represent living birds. As such, our experiment did not take account of the many additional parameters that may affect detection of living birds, the majority of which survive stranding (Rodríguez et al. 2017) and likely move away from their original stranding locations (Telfer et al. 1987; Rodríguez et al. 2017; pers. obs.). These parameters should thus be estimated and incorporated into seabird stranding data correction factors: namely, spatial fallout patterns, post-stranding movement patterns, and predation rates of living stranded seabirds; persistence and reporting rates of carcasses; and rescuer search patterns. For example, spatial fallout patterns could be better discerned by radio- or GPS-tagging fledglings before they depart the colony and tracking their movements (e.g., Rodríguez et al. 2022), or by conducting standardized searches across all light and dark areas in the potential fallout zone (as suggested in Brown et al. 2023). The fates of individuals that remain undetected by rescuers are largely unknown but could be studied by tracking movements and predation of stranded seabirds using cameras mounted at stranding hotspots, thermal imaging drones, or manually tracking stranded birds that have been fitted with radio-frequency tags prior to fledging (e.g., in small populations). Meanwhile, rescuer search patterns, which may be biased toward illuminated areas (suggested by Brown et al. 2023, this study), could be tracked across the entire search area using individual GPS units and subsequent characterization of the light levels and calculation of proportions of time spent searching in each sector of the search area. Collectively, better understanding of such parameters would allow more precise estimates of seabird detection rate by rescuers and therefore also of the true magnitude of stranding-induced mortality.

## Conclusion

Using our approximations of detection probability calculated using randomly-placed, stationary, plastic decoys, we estimate that at least 10% of stranded pufflings go undetected by Puffin Patrol rescuers at hotspot stranding sites. In areas of fallout zones that receive lower search effort, in locations that are dark and intermediately concealed, and in smaller-bodied seabirds, the proportion of undetected individuals may be much higher. Our study confirms that mortality associated with strandings may be severely underestimated (Rodríguez et al. 2017), with potentially grave consequences for seabird populations already in decline, and offers a first step in the development of correction factors which can be adapted to site- and species-specific requirements of other rescue programs around the world that rely on volunteers to search for stranded birds.

## Acknowledgments

We thank the Canadian Parks and Wilderness Society – Newfoundland and Labrador Chapter, Puffin Patrol staff, and all the volunteer Puffin Patrollers for their logistical assistance and participation in the study; Michael Campbell for early inspiration and idea formation; Lincoln Savi from SaviMade for the high-quality 3D puffin decoy file design and finished product; Dave Fifield for help with 3D-printing; Wesley Burr for help and advice in early experimental design; Madeline Sceviour and Fiona Le Taro for help with fieldwork and equipment; Laura McKinnon and Joshua Wilson for analysis assistance and advice; Liana Brown for guidance in human research ethics; Hani at “Grocery Deli and More” convenience store as well as the various managers and workers at the BGI crab plant for their hospitality and cooperation.

## Ethics statement

All procedures were performed in accordance with Trent University’s Research Ethics Board guidelines under protocol #28055. We created an informed consent agreement which was approved by the Trent University Research Ethics Board before the experiment (see Supplemental Material). As part of the stipulations of the protocol, we as the “Experimenters” (T.M.B. and L.v.H.) were not in contact with study participants during the signup process, were not privy to their identities or information, and were not allowed to ask their participatory status in the experiment when we encountered them at the study sites.

Puffin Patrol volunteers who arrived at the headquarters at 2200 hours on the nights of this experiment were read a short script by program organizers, requesting their participation in this experiment. Those who consented to participate were administered a consent agreement (see Supplemental Material). A total of *N* = 55 agreements were signed by Puffin Patrol volunteers who chose to participate in the study (i.e., to collect decoys). Puffin Patrol organizers showed volunteers a sample decoy to familiarize them with its appearance, and specifically asked volunteers who decided not to participate to leave decoys in place whenever they encountered them. Participants were not told how many decoys would be deployed, so as not to influence their search effort. Participants who participated on two or more nights of the experiment were not required to sign new consent agreements. As such, any repeat Participants are not necessarily accounted for in the *N* = 55 consent agreements collected and it is therefore unknown exactly how many Participants were involved in this experiment. Personal information of Participants was kept separate from any experimental data to maintain anonymity.

## Author contributions

T.M.B. conceived the ideas and formulated hypotheses, designed the field protocols and methods, collected and analyzed data, and wrote the first version of the manuscript. L.v.H. helped design field protocols and collect data. S.I.W. helped supervise the research and edited the manuscript. S.D. helped design the field protocols and methods and edited the manuscript. G.B. supervised the research, helped design field protocols and methods, and substantially edited the manuscript.

## Data availability

Data will be available on Dryad.

