## Supplemental Material for "Artificial light at night and concealment level affect detection probability of stranded seabirds: a study with 3D-printed puffin decoys"

Methods

*Materials*

Decoys were constructed of black polylactic acid plastic using a filament-based 3D printer (Creality CR10S; Shenzhen, China). Right and left legs were designed and printed separately, then inserted and glued into sockets in the main body of the decoy. We spray-painted decoy undersides white in a pattern similar to that of real pufflings (Krylon Fusion All-In-One Paint and Primer in Gloss Dover White; Sherwin-Williams; Cleveland, OH, USA) and used black permanent marker to write a unique identifying number (1-60) on the underside of the tail.

*Participants*

<u>Pre-experiment script to be read to volunteers</u>

"This year, the Puffin Patrol is working with a PhD student at Trent University named Taylor Brown. Taylor is studying puffins and why they get stranded here in coastal Newfoundland. Taylor is running an experiment with our help, and we are wondering if you would like to participate in it. It is completely optional. Taylor has put some fake, plastic puffins like this [show them the puffin model] in random places where you will be searching for real puffins. If you want to participate in this experiment, all you have to do is sign this consent form and bring back to us any plastic puffins that you find while you are out looking for real ones. Alternatively, you can give the plastic puffins to Taylor or assistant Lucy, who will be present in the area with vests on that say their names. Your name and signature on the form will be kept separate from any data you help collect, so you will stay totally anonymous. If you sign the consent form but change your mind about participating, just let us know by the end of the night."

Consent Agreement

The formal consent agreement for this study is on file with Trent University's Research Ethics Board under protocol #28055.

*Statistical Analysis*

Decoy detection rate relative to levels of illumination, concealment, and search effort

To complement our statistical analysis, we modeled persistence probability of puffling decoys relative to each level of our three covariates of interest (light level, concealment level, and search effort). Persistence probability is the probability of decoys remaining undiscovered over time, whereby a high persistence probability generally indicates a low detection rate (i.e., it took longer for Participants to find the decoys) and vice versa. The aim was to display how long (in elapsed time) real pufflings may await rescue. Because search effort was a continuous variable measured in search-minutes, we created three arbitrary bins (i.e., categories) for better visualization. These bins of search effort were "low" (≤60 search-minutes), "medium" (61-120 search-minutes), and "high" (≥121 search-minutes). Across persistence probability models for all three covariates of interest, we modeled only the main effect, with no random effects, stratified variables, or determinants of significance, again because these models were created for visualization purposes of the main effects only. It is important to note that the main way in which modeled persistence probabilities visually reflect significant differences in detection rate is in the absence of model intersections and overlaps; lack of differences in detection rate are visualized as intersecting and overlapping persistence probability models throughout the duration of the experiment. Confidence intervals should therefore be interpreted with caution.

*Results*

Decoy Detection Rate Relative to Levels of Illumination, Concealment, and Search Effort

(i) Persistence probability of decoys in relation to light levels

As was the case in our formal detection rate analysis (i.e., Cox proportional hazards model), decoys deployed in bright light trended toward having a lower probability of persistence (i.e., were more easily found) over the course of the experiment than those in the dark, but models intersected several times during the first half of the experiment (Figure S2). Meanwhile, puffling decoys in dim light had intermediate persistence probability. Our models demonstrate that if a decoy had not been found within approximately 90 minutes from the beginning of the experiment, it was unlikely to be found, irrespective of lighting conditions (Figure S2).

(ii) Persistence probability of decoys in relation to concealment levels

Persistence probability decreased steadily but irregularly across the four concealment levels, with several equivalence points (i.e., model crossovers) in the first 40 minutes of the experiment (Figure S3). After 90 minutes, few of the remaining decoys were found and persistence probability leveled off at approximately 0.2 for decoys in the open and those concealed on three sides, approximately 0.3 for decoys concealed on one side, and approximately 0.4 for decoys concealed on two sides (Figure S3).

(iii) Persistence probability in relation to search effort

Persistence probabilities decreased over time but did not vary among decoys subjected to different levels of search effort (Figure S4). Although detection rate increased by 0.6% per minute of expended search effort, the only visible trend in the plotted persistence probabilities was that persistence was higher when a site was subjected to low search effort, compared to medium or high (Figure S4).

*Tables*

**Supplemental Material Table S1.** All decoy locations at both of the experimental sites, including their coordinates (easting and northing), illuminance (lux), qualitative light level, and concealment level.

| Site | Location | Easting | Northing | Illuminance (lux) | Light Level | Concealment Level |
|---|---|---|---|---|---|---|
| Store | B01 | 361486 | 5237789 | 0 | Dim | 0 |
| Store | B02 | 361477 | 5237808 | 30 | Bright | 2 |
| Store | B03 | 361513 | 5237828 | 0 | Dim | 3 |
| Store | B04 | 361493 | 5237810 | 0 | Dim | 0 |
| Store | B05 | 361514 | 5237838 | 14 | Bright | 0 |
| Store | B06 | 361510 | 5237822 | 9 | Dim | 0 |
| Store | B07 | 361501 | 5237834 | 0 | Dim | 0 |
| Store | B08 | 361496 | 5237826 | 29 | Bright | 1 |
| Store | B09 | 361487 | 5237814 | 0 | Dim | 0 |
| Store | B10 | 361476 | 5237831 | 19 | Bright | 0 |
| Store | B11 | 361477 | 5237816 | 13 | Bright | 0 |
| Store | B12 | 361509 | 5237832 | 14 | Bright | 1 |
| Store | B13 | 361507 | 5237823 | 0 | Dim | 0 |
| Store | B14 | 361477 | 5237798 | 0 | Dim | 0 |
| Store | B15 | 361480 | 5237824 | 0 | Dim | 2 |
| Store | B16 | 361483 | 5237818 | 0 | Dark | 2 |
| Store | B17 | 361508 | 5237769 | 0 | Dark | 0 |

| | | | | | | |
|---|---|---|---|---|---|---|
| Store | B18 | 361500 | 5237765 | 0 | Dark | 0 |
| Store | B19 | 361483 | 5237767 | 0 | Dark | 1 |
| Store | B20 | 361494 | 5237748 | 0 | Dark | 0 |
| Store | B21 | 361494 | 5237745 | 0 | Dark | 2 |
| Store | B22 | 361525 | 5237811 | 0 | Dark | 0 |
| Store | B23 | 361502 | 5237755 | 0 | Dark | 0 |
| Store | B24 | 361504 | 5237781 | 0 | Dark | 0 |
| Store | B25 | 361513 | 5237810 | 0 | Dark | 1 |
| Store | B26 | 361479 | 5237784 | 0 | Dark | 1 |
| Store | B27 | 361530 | 5237826 | 0 | Dark | 1 |
| Store | B28 | 361492 | 5237834 | 0 | Dark | 1 |
| Store | B29 | 361499 | 5237814 | 0 | Dark | 3 |
| Store | B30 | 361480 | 5237842 | 0 | Dark | 3 |
| Plant | C01 | 362100 | 5237785 | 10 | Bright | 1 |
| Plant | C02 | 362111 | 5237797 | 13 | Bright | 2 |
| Plant | C03 | 362099 | 5237802 | 66 | Bright | 0 |
| Plant | C04 | 362112 | 5237803 | 16 | Bright | 0 |
| Plant | C05 | 362118 | 5237800 | 0 | Dim | 1 |
| Plant | C06 | 362120 | 5237832 | 32 | Bright | 0 |
| Plant | C07 | 362116 | 5237838 | 46 | Bright | 1 |
| Plant | C08 | 362095 | 5237839 | 24 | Bright | 0 |
| Plant | C09 | 362112 | 5237845 | 35 | Bright | 3 |
| Plant | C10 | 362110 | 5237852 | 21 | Bright | 0 |
| Plant | C11 | 362096 | 5237856 | 21 | Bright | 1 |
| Plant | C12 | 362066 | 5237866 | 10 | Bright | 1 |

| | | | | | | |
|---|---|---|---|---|---|---|
| Plant | C13 | 362063 | 5237850 | 38 | Bright | 3 |
| Plant | C14 | 362042 | 5237845 | 42 | Bright | 1 |
| Plant | C15 | 362079 | 5237764 | 10 | Bright | 3 |
| Plant | C16 | 362069 | 5237736 | 0 | Dark | 1 |
| Plant | C17 | 362131 | 5237804 | 0 | Dark | 1 |
| Plant | C18 | 362136 | 5237819 | 0 | Dark | 0 |
| Plant | C19 | 362131 | 5237831 | 0 | Dark | 3 |
| Plant | C20 | 362122 | 5237862 | 0 | Dark | 1 |
| Plant | C21 | 362090 | 5237861 | 0 | Dark | 1 |
| Plant | C22 | 362089 | 5237878 | 0 | Dark | 2 |
| Plant | C23 | 362082 | 5237862 | 0 | Dark | 3 |
| Plant | C24 | 362084 | 5237855 | 0 | Dark | 1 |
| Plant | C25 | 362047 | 5237845 | 0 | Dark | 2 |
| Plant | C26 | 362035 | 5237841 | 0 | Dark | 1 |
| Plant | C27 | 362040 | 5237842 | 0 | Dark | 2 |
| Plant | C28 | 362109 | 5237869 | 0 | Dark | 2 |
| Plant | C29 | 362093 | 5237825 | 0 | Dark | 2 |
| Plant | C30 | 362090 | 5237779 | 0 | Dark | 4 |

*Figures*

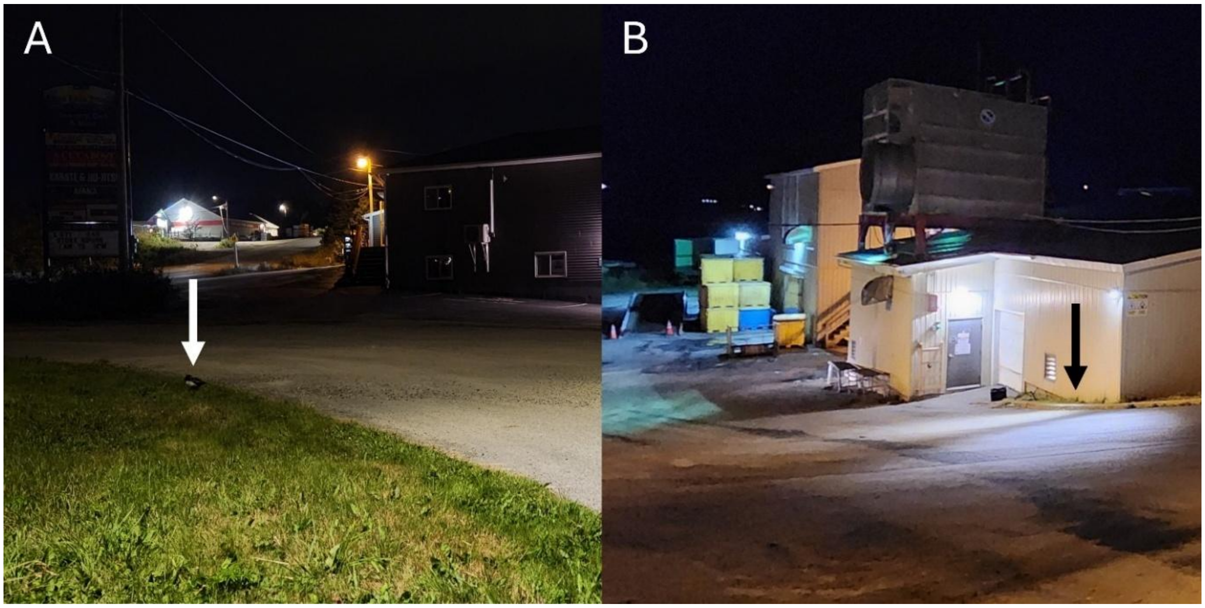


**Supplemental Material Figure S1.** A) Example of a "dim" illuminated location with concealment score of 0 at the convenience store site. A puffling decoy is indicated by white arrow. B) Example of a “bright” location with concealment score of 1 (indicated by black arrow) at the crab plant site (note that a decoy is not present in picture).

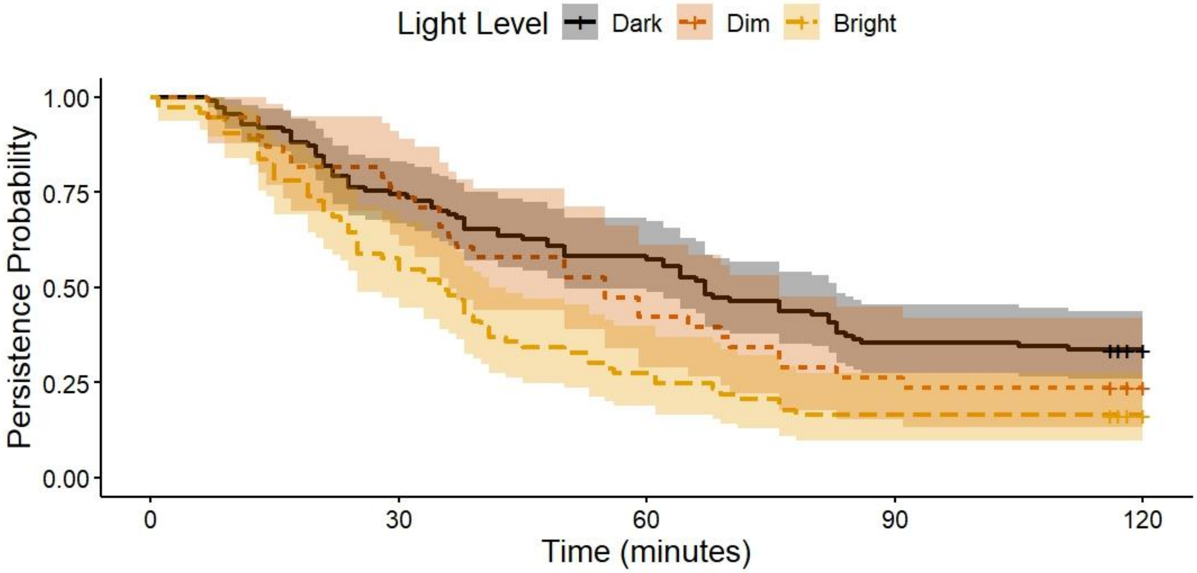


**Supplemental Material Figure S2.** Light level persistence curves, depicting main effects for visualization purposes only. Persistence probability of decoys (as a proportion of 1.00) over the duration of the experiment (in minutes) deployed in three different light level conditions: darkness, dim illumination, and bright illumination. Persistence probability is the probability of decoys remaining undiscovered over time, whereby a high persistence probability indicates a low detection rate over time. Censored individuals are indicated on the figure by “+”, which in these cases were deployed decoys that persisted the entire experiment, but the experiment duration was slightly less than 120 minutes (*N* = 22 occasions). Shaded areas represent 95% confidence intervals.

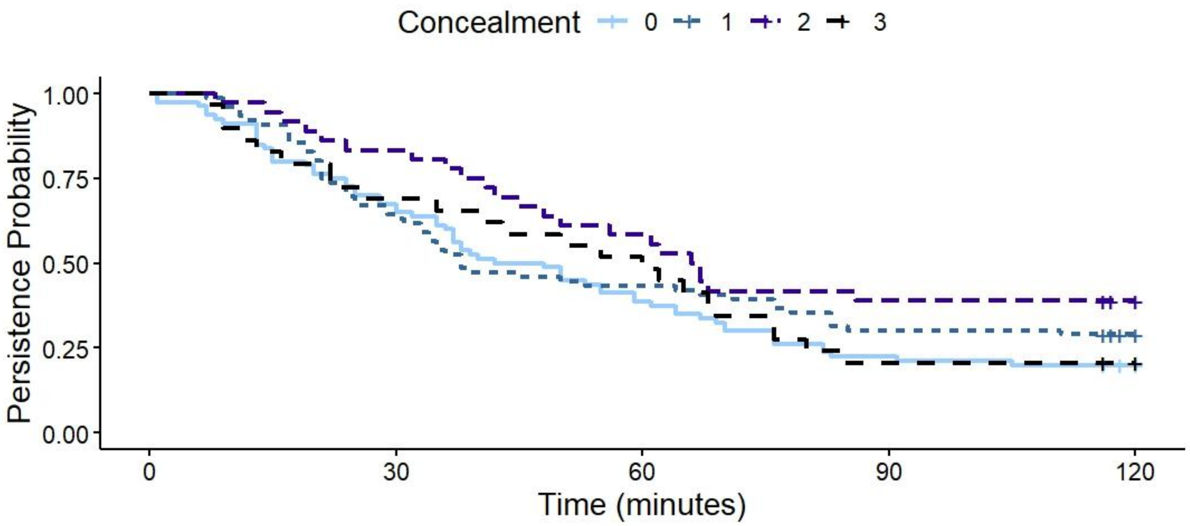


**Supplemental Material Figure S3.** Concealment level persistence curves, depicting main effects for visualization purposes only. Persistence probability of decoys (as a proportion of 1.00) over the duration of the experiment (in minutes) deployed in four different concealment levels: completely unconcealed (score of 0), and concealed on one, two, and three sides. See Figure S2 for an explanation of persistence probability and details of censored individuals. Confidence intervals overlapped across concealment levels and were removed for readability.

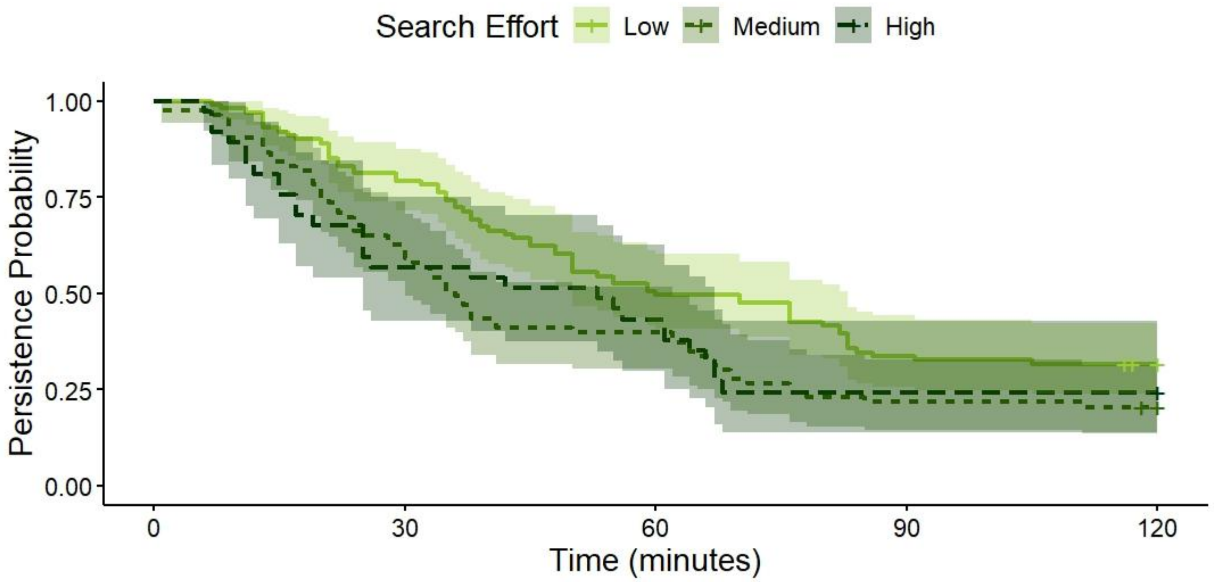


**Supplemental Material Figure S4.** Search effort persistence curves, depicting main effects for visualization purposes only. Persistence probability of decoys (as a proportion of 1.00) over the duration of the experiment (in minutes) when exposed to different levels of search effort. Search effort was statistically analyzed as a continuous variable, but for visualization purposes we created three bins: low (≤60 search-minutes; $N = 101$), medium (61-120 search-minutes; $N = 83$), and high (≥121 minutes; $N = 37$) search effort. See Figure S2 for an explanation of persistence probability and details of censored individuals. Shaded areas represent 95% confidence intervals.
